# Impact of NaCl, nitrate and temperature on the microbial community of a methanol-fed, denitrifying marine biofilm

**DOI:** 10.1101/607028

**Authors:** Richard Villemur, Geneviève Payette, Valérie Geoffroy, Florian Mauffrey, Christine Martineau

## Abstract

**Background:** The biofilm of a continuous, methanol-fed, fluidized denitrification system that treated a marine effluent at the Montreal Biodome is composed of a multi-species microbial community, among which *Hyphomicrobium nitrativorans* NL23 and *Methylophaga nitratireducenticrescens* JAM1 are the principal bacteria involved in the denitrifying activities. To assess its resilience to environmental changes, the biofilm taken from the denitrification system was cultured at laboratory scale in artificial seawater (ASW) under anoxic conditions and exposed to a range of specific physico-chemical parameters. We previously showed that the seawater formulation and the NaCl concentrations had a strong impact on the *H. nitrativorans* NL23 population, with its displacement by a new denitrifier, *M. nitratireducenticrescens* GP59. Here, we report the impact of these cultures conditions on the dynamics of the overall microbial community of the denitrifying biofilm.

**Methods:** The original biofilm (OB) taken from the denitrification system was acclimated for five weeks in ASW under anoxic conditions with a range of NaCl concentrations, and with four combinations of nitrate concentrations and temperatures. The OB was also acclimated to the commercial Instant Ocean seawater medium (IO). The bacterial diversity of the biofilm cultures and the OB was determined by 16S ribosomal RNA amplicon metagenome sequencing. Culture-dependent approach was used to isolate other denitrifying bacteria from the biofilm cultures. The metatranscriptomes of some of the biofilm cultures were derived, along with the transcriptomes of planktonic pure cultures of *H. nitrativorans* NL23 and *M. nitratireducentricrescens* GP59 cultivated under denitrifying conditions.

**Results:** The 16S metagenomic data revealed very high proportions of *M. nitratireducenticrescens* in the biofilm cultures. *H. nitrativorans* NL23 was found in high proportion in the OB, both was absent in the biofilm cultures with 2.75% NaCl in the ASW medium. It was found however in low proportions in the biofilm cultures with 0 to 1% NaCl in the ASW medium and in the IO biofilm cultures. Emergence of *Marinicella* spp. occurred in these biofilm cultures. Denitrifying bacterial isolates affiliated to *Marinobacter* spp. and *Paracoccus* spp. were isolated. Up regulation of the denitrification genes in strains GP59 and NL23 occurred in the biofilm cultures compared to the planktonic pure cultures. Denitrifying bacteria affiliated to the *Stappia* spp. were metabolically active in the biofilm cultures.

**Conclusions:** These results illustrate the dynamics of the microbial community in the denitrifying biofilm cultures in adapting to different environmental conditions. The NaCl concentration is an important factor affecting the microbial community in the biofilm cultures. Up regulation of the denitrification genes in strain GP59 and strain NL23 in the biofilm cultures suggests different mechanisms of regulation of the denitrification pathway in the biofilm compared to the planktonic pure cultures. Other denitrifying heterotrophic bacteria are present in low proportions in the biofilm, suggesting that the biofilm has the potential to adapt to heterotrophic, non-methylotrophic environments.

## Introduction

Most naturally-occurring microbial biofilms, such as those encountered in bioremediation processes, are composed of multiple microbial species. Studying such complex biofilms is a challenge, as each species can influence the biofilm development. The biofilm microbial community inside a bioremediation process adapts to the prescribed operating conditions and shapes the efficiency of the bioprocess to degrade the pollutant(s). Usually, the microbial community in such bioprocesses is complex and composed of main degraders but also of secondary microorganisms that could provide benefits to the degraders or could simply contribute to the degradation intermediates or waste. It is recognized that a complex microbial community is more resilient to “unexpected” changes in the operation of the bioprocesses than single species biofilm, as some of the minor degraders take over the main degraders affected by the changes (Cabrol and Malhautier, 2011; Salta *et al.*, 2013; Roder *et al.*, 2016; Tan *et al.*, 2017). The mechanisms of how a microbial population in a biofilm adapts to changes are however poorly understood.

The Montreal Biodome, a natural science museum, operated a continuous fluidized methanol-fed denitrification reactor to remove nitrate (NO_3_^−^) that accumulated in the 3 million-L seawater aquarium. The fluidized carriers in the denitrification reactor were colonized by naturally-occurring multispecies microorganisms to generate a marine methylotrophic denitrifying biofilm estimated to be composed by around 15-20 bacterial species. The main bacteria responsible of the denitrifying activities belong to the alphaproteobacteria *Hyphomicrobium nitrativorans* (strain representative NL23) and to the gammaproteobacteria *Methylophaga nitratireducenticrescens* (strain representative JAM1), both methylotrophs, that accounted for 60-80% of the biofilm (Labbé *et al.*, 2003; Labbé *et al.*, 2007; Martineau *et al.*, 2013b; *Villeneuve et al., 2013*).

Denitrification takes place in bacterial cells where N oxides serve as terminal electron acceptor instead of oxygen (O_2_) for energy production when oxygen depletion occurs, leading to the production of gaseous nitrogen (N_2_). Four sequential reactions are strictly required for the reduction of NO_3_^−^ to gaseous nitrogen, via nitrite (NO_2_^−^), nitric oxide (NO) and nitrous oxide (N_2_O), and each of these reactions is catalyzed by different enzymes, namely NO_3_^−^ reductases (Nar and Nap), NO_2_^−^ reductases (NirS and NirK), NO reductases (Nor) and N_2_O reductases (Nos) (Philippot and Hojberg, 1999; Richardson *et al.*, 2001; Kraft *et al.*, 2011). Whereas *H. nitrativorans* NL23 possesses the four reductases for the complete denitrification pathway, *M. nitratireducenticrescens* JAM1 performs incomplete denitrifying activities, as it lacks a dissimilatory NO-forming nitrite reductase (Auclair *et al.*, 2010; Villeneuve *et al.*, 2012; Martineau *et al.*, 2013a; Mauffrey *et al.*, 2015; Mauffrey *et al.*, 2017). Using degenerated PCR primers for the detection of denitrification genes, we showed that there are probably other denitrifying bacteria in the biofilm, one to four orders of magnitude lower in concentrations than *M. nitratireducenticrescens* JAM1 and *H. nitrativorans* NL23 (Auclair *et al.*, 2012). These other bacteria may play a role if the bioprocess underwent stress conditions or changes in the operation mode.

Because of the relatively low number of bacterial species, this denitrifying biofilm offers an excellent model to study the evolution and the dynamics of a naturally-occurring microbial community in a biofilm. We have initiated a study to assess the performance of the denitrifying biofilm subjected to different conditions. In the first two parts of the study reported by Geoffroy *et al.* (2018) and Payette *et al.* (2019), the original biofilm (OB) taken from the Biodome denitrification system was cultured in an artificial seawater (ASW) under batch-mode, anoxic conditions at laboratory scale and exposed to a range of specific physico-chemical parameters. Such parameters included a range of NaCl and NO_3_^−^ concentrations, and of temperatures. We showed that the seawater formulation and the NaCl concentrations have significant impacts on the *H. nitrativorans* NL23 population, with its displacement by a subpopulation of the species *M. nitratireducenticrescens* (strain GP59 as a representative), which can perform the complete denitrification pathway. In the last part of this study, reported here, we measured the impact of these culture conditions on the dynamics of the overall microbial community of the denitrifying biofilm. The 16S ribosomal RNA (rRNA) gene sequences were derived from these biofilm cultures to compare the evolution of the bacterial community between culture conditions. The metatranscriptomes of three biofilm cultures were also derived to assess changes in metabolic pathways of *H. nitrativorans* NL23 and *M. nitratireducentricrescens* GP59 between the planktonic pure cultures and the biofilm cultures. These metatranscriptomes were also analyzed to assess the metabolic contributions of other microorganisms in the biofilm cultures. Moreover, by using culture dependent approaches, denitrifying bacterial isolates affiliated to *Paracoccus* spp. and *Marinobacter* spp. were recovered from the biofilm cultures. Our results showed that changes in the expression of denitrification genes occurred between the planktonic pure cultures and the biofilm cultures for *M. nitratireducentricrescens* GP59 and *H. nitrativorans* NL23. Finally, metatranscriptome analyses revealed the presence of active denitrifying bacteria affiliated to the genus *Stappia* in the biofilm cultures. Our results demonstrated the dynamics and the plasticity of the microbial community in the denitrifying biofilm in adapting to different environmental conditions.

## Material and Methods

### Acclimation of the original biofilm to different physico-chemical parameters

The formulations of the artificial seawater (ASW) medium and the commercial Instant Ocean (IO) medium, and the different conditions of the biofilm cultures were described by Payette *et al.* (2019). Briefly, the biomass of several carriers taken from the denitrification reactor of the Montreal Biodome was scrapped, dispersed, then distributed to several vials containing twenty free carriers and 60 mL prescribed medium (Table 1; Fig. 1). The vials were incubated under anoxic conditions at 23°C or 30°C (Table 1) and shaken at 100 rpm (orbital shaker). In average once a week, the twenty carriers were taken, gently washed to remove the excess medium and the planktonic bacteria, then transferred into fresh anoxic medium and incubated under the same conditions (Fig. 1). The Ref300N-23C biofilm cultures (for 300 mg NO3--N/L, 23°C) were defined as the *Reference biofilm cultures.* These cultures were used by Payette *et al.* (2019) as a reference to compare results between the different culture conditions. The protocols to measure NO_3_^−^ and NO_2_^−^ concentrations, and to extract DNA from the biofilm cultures or the planktonic pure cultures were described in Payette *et al.* (2019) and Geoffroy *et al.* (2018).

**Figure 1.**
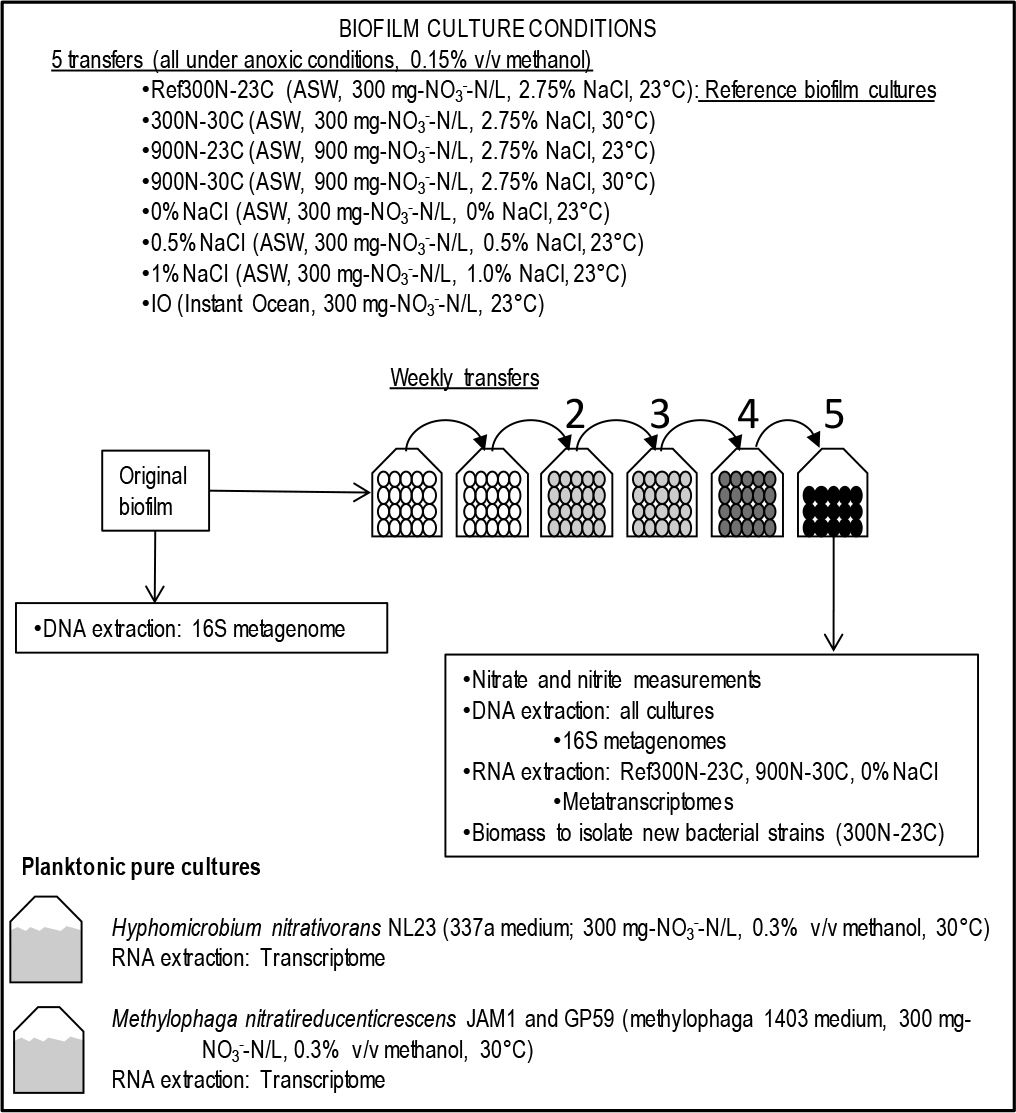
Schematic of the experimental assays. The original biofilm was thawed, scrapped from the carriers, dispersed and distributed in vials containing the ASW medium supplemented with prescribed concentrations of NO3−, methanol and NaCl, or containing the IO medium (Table 1), and 20 free carriers. The vials were incubated at the prescribed temperature under anoxic conditions. The carriers were transferred five times at appr. 1-week interval into fresh prescribed medium and incubated in the same conditions. NO3− and NO2− concentrations were measured every 24-48 h. Methanol and NaNO3 were added when needed if NO3− was completely depleted between transfers. DNA or RNA was extracted from the biofilm at the end of the 5th transfer cultures. Planktonic pure cultures were carried out in recommended culture medium for the respective strains.

**Table 1.** Biofilm culture conditions

| Name | Medium | NO<sub>3</sub><sup>-</sup> | Methanol | NaCl | Temp | Denitrif<sup>a</sup> rates |
| --- | --- | --- | --- | --- | --- | --- |
|  |  | mM | % (v/v) | % (w/v) | °C | mM-NO<sub>x</sub> h<sup>-1</sup> vial<sup>-1</sup> |
| Ref300N-23C<sup>b</sup> | ASW | 21.4 (300) | 0.15 | 2.75 | 23 | 1.47 |
| 300N-30C | ASW | 21.4 (300) | 0.15 | 2.75 | 30 | 1.94 |
| 900N-23C | ASW | 64.3 (900) | 0.45 | 2.75 | 23 | 1.95 |
| 900N-30C | ASW | 64.3 (900) | 0.45 | 2.75 | 30 | 2.37 |
| 0% NaCl | ASW | 21.4 (300) | 0.15 | 0 | 23 | 1.28 |
| 0.5% NaCl | ASW | 21.4 (300) | 0.15 | 0.5 | 23 | 1.57 |
| 1% NaCl | ASW | 21.4 (300) | 0.15 | 1.0 | 23 | 0.66 |
| IO | IO | 21.4 (300) | 0.15 | 3.0<sup>c</sup> | 23 | 0.76 |

The original biofilm was cultured in triplicates in these conditions at pH 8.0. The carriers were transferred 5 times in fresh medium around each week before measuring the denitrifying activities.
<sup>a</sup> From Payette *et al.* (2019)
<sup>b</sup> Reference cultures.
<sup>c</sup> The exact amount of NaCl added in the IO medium is not known. See Payette *et al.* (2019) for the IO composition. For comparison, the amount of Na<sup>+</sup> and Cl<sup>-</sup> in the ASW medium is 3.2%.
In gray are changed parameters from the reference biofilm cultures.
IO: Instant Ocean medium.

### Metagenomes of 16S rRNA gene sequences

DNA extracted from triplicate biofilm cultures was pooled before sequencing. Total DNA samples from seven biofilm cultures (Table 1; Ref300N-23C, 300N-30C, 900N-23C, 900N-30C, 0%NaCl, 0.5%NaCl and 1.0%NaCl) were sent to the sequencing service at the Research and Testing Laboratory (Lubbock, Texas, USA). A region of the 16S rRNA genes was PCR amplified using the 28F-519R primers (5’ GAGTTTGATCNTGGCTCAG 3’ and 5’ GTNTTACNGCGGCKGCTG 3’, covering the V1-V2-V3 variable regions) and subjected to pyrosequencing using a Roche 454 FLX genome sequencer system. Between 8562 and 14048 high quality reads with an average of 450 nt were obtained and analyzed to remove chimera (UCHIME). Samples from the original biofilm [OB] and the IO biofilm cultures (Table 1) were sent two years later to the sequencing service of Genome Quebec Innovation Center (Montreal, QC, Canada). In these cases, the 16S rRNA sequences covering the V6-V7-V8 variable regions (5’ ACACTGACGACATGGTTCTACA 3’ and 5’ TACGGTAGCAGAGACTTGGTCT 3’) were PCR amplified and sequenced by Illumina MiSeq PE250 (paired-end). The paired-end high quality reads were assembled resulting in app. 420-nt sequences. Chimeric sequences were removed by UCHIME. These resulted in 346158 sequences for the OB sample and 317977 sequences for the IO sample. Sequences were then clustered with 3% divergence. All representative sequences of the OTUs (from pyrosequencing and Illumina) were checked again for chimeras with the DECIPHER v 2.0 program (http://www2.decipher.codes/FindChimeras.html) (Wright *et al.*, 2012). The affiliation of the OTUs to the most probable genus was determined by the CLASSIFIER program at the Ribosomal data project (RDP) web site (Cole *et al.*, 2014). 16S rRNA sequence reads were deposited in the GenBank Sequence Read Archive (SRA) under the accession number PRJNA524642.

### Isolation of bacterial isolates

Biofilm of the Ref300C-23C biofilm cultures was scrapped from the carriers and dispersed in saline solution (3% NaCl, 34.2 mM phosphate buffer pH 7.4), and serial dilutions were made and inoculated onto these agar plate media: (1) R2A medium (complex organic carbons; EMD Chemicals Inc., Gibbstown, NJ, USA), (2) Marine Agar 2216 (marine medium with yeast extract and peptone as carbon source; Becton, Dickinson and Co., Sparks, MD, USA), (3) methylophaga medium 1403 (American Type Culture Collection [ATCC], Manassas, VA, USA) and (4) the ASW medium; these two latter media were supplemented with 1.5% agar and 0.3% v/v methanol. The isolation procedure, the taxonomic affiliation of the isolates and the measurement of their denitrifying activities were carried out as described by Geoffroy *et al.* (2018). The 16S rRNA gene sequences were deposited in GenBank under the accession numbers MK571459 to MK571476.

### Transcriptomes

Planktonic pure cultures of *M. nitratireducenticrescens* strains JAM1 and GP59 were performed in the methylophaga 1403 medium and of *H. nitrativorans* strain NL23 in the 337a medium as described by Martineau *et al.* (2015) and Mauffrey *et al.* (2015). These cultures were carried out in triplicate with methanol (0.3%) and NO_3_^−^ (21.4 mM [300 mg-N/L]) under anoxic conditions at 30°C. The biomass was collected by centrifugation when the NO_3_^−^ reduction was near completion, and total RNA was extracted as described by Mauffrey *et al.* (2015). For the biofilm cultures, at the end of the acclimation assays (the fifth transfer), the biomass of each replicate was scrapped from carriers and used to extract total RNA. The RNA samples were sent to the sequencing service for RNA sequencing (RNAseq) by Illumina (Genome Quebec Innovation Center, Montreal QC, Canada). Because of limited amount of biofilm available, total RNA from the triplicate biofilm samples were pooled before sending to the sequencing service. For the planktonic pure cultures, RNAseq was performed on each replicate. The Ribo-Zero™ rRNA Removal Kit (Meta-Bacteria; Epicentre, Madison, WI, USA) was used to deplete total RNA of the ribosomal RNA. The RNA was then treated with the TruSeq Stranded mRNA Sample Prep Kit (Illumina Inc, San Diego, CA, USA).

All computations were made on the supercomputer Briarée from the Université de Montréal, managed by Calcul Québec and Compute Canada. Raw reads were filtered to remove low quality reads using FASTX toolkit (http://hannonlab.cshl.edu/fastx_toolkit/) by discarding any reads with more than 10% nucleotides with a PHRED score <20. The resulting reads from each sample/replicate were aligned respectively to the genome of *M. nitratireducenticrescens* JAM1 (GenBank accession number CP003390.3), to the genome and plasmids of *M. nitratireducenticrescens* GP59 (CP021973.1, CP021974.1, CP021975.1) and to the genome of *H. nitrativorans* NL23 (CP006912.1) using Bowtie (v 2.2.3) with default parameters. SAMtools (v 0.1.18) and BEDtools (v 2.20.1) were used for the generation of sam and bam files, respectively. Significance for difference in the relative transcript levels of the corresponding genes/sequences (defined as transcript per million: TPM) between strain JAM1 (average of triplicate planktonic cultures), strain GP59 (average of duplicate planktonic cultures), strain NL23 (average of triplicate planktonic cultures) and the respective biofilm cultures was performed with the R Bioconductor NOIseq package v2.14.0 (NOIseqBio) (Tarazona *et al.*, 2011) and run with the R software v3.2.3 (Team, 2015). Because the RNAseq from the biofilm samples were derived from one pooled RNA preparation, the “no replicate parameter” was set (*pnr*=0.2, *nss*=5 and *v*=0.02; pseudoreplicate generated) in NOIseq. Results from this statistical analysis showed that genes/sequences that had at least >2-fold higher transcript levels from one type of culture to the other showed significant differences. RNAseq reads from the planktonic pure cultures and the biofilm cultures were deposited in the SRA under the accession number PRJNA525230. Annotations were based on services provided by GenBank (https://www.ncbi.nlm.nih.gov/genbank), RAST (Rapid Annotation using Subsystem Technology; http://rast.nmpdr.org) and KEGG (Kyoto Encyclopedia of Genes and Genomes; https://www.genome.jp/kegg).

To derive transcript reads not associated to the *M. nitratireducenticrescens* and *H. nitrativorans*, the filtered, high quality reads were aligned to a concatenated sequence consisting of the three reference genomes (JAM1+GP59+NL23) and the two plasmids (from strain GP59). The reads that did not align were kept. The FastQ files of the unaligned reads were *de novo* assembled at the National Center for Genome Analysis web site (https://galaxy.ncgas-trinity.indiana.edu) by Trinity v. 2.4.0 (Grabherr *et al.*, 2011). These transcripts were deposited in SRA under the accession number PRJNA525230. Estimation of the transcript abundance of the *de novo* assembled sequences was performed by RSEM (Li and Dewey, 2011). The resulting assembled sequences were annotated at the Joint Genomic Institute (https://img.jgi.doe.gov/cgi-bin/m/main.cgi) to find open reading frames with their putative function and affiliation (GOLD Analysis Project Id: Ga0307915, Ga0307877, Ga0307760). The annotations were then verified manually for discrepancies within the assembled sequences.

## Results

### Bacterial composition of the biofilm cultures by 16S metagenomic analysis

As reported by Payette *et al.* (2019), the original biofilm (OB) collected from the Biodome denitrification system was used as inoculum in a series of biofilm cultures cultivated under anoxic conditions in an homemade artificial seawater (ASW) medium. These biofilm cultures were acclimated to different conditions that include a range of NaCl concentrations from 0% to 2.75%, four combinations of NO_3_^−^ concentrations (300 and 900 NO_3_^−^-N/L) and temperatures (23 and 30°C) (Table 1; Fig. 1). In addition, the biofilm was acclimated to the commercial Instant Ocean (IO) medium used by the Biodome. The composition of the bacterial community of these acclimated biofilm cultures was determined by sequencing of the 16S rRNA genes (16S) to assess the evolution of the bacteria community under the specific acclimated conditions (Fig. 2; Table 2).

**Figure 2.**
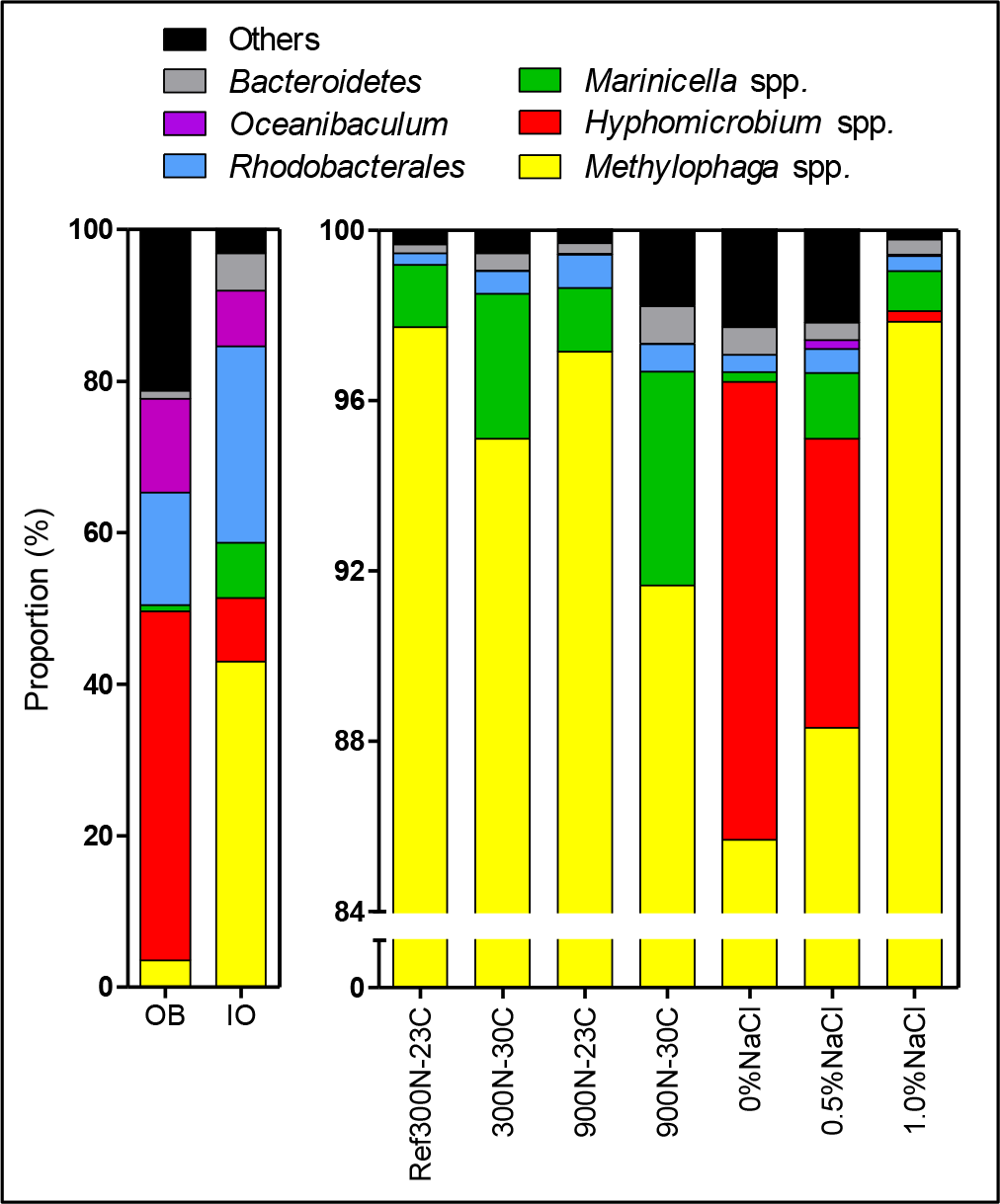
Proportion of affiliated OTUs in the biofilm cultures. Bacterial composition of OB and the IO biofilm cultures was determined by sequencing the V6-V7-V8 variable regions of the 16S rRNA gene by Illumina, whereas the other samples were determined by sequencing the V1-V2-V3 variable regions by pyrosequencing.

**Table 2.**
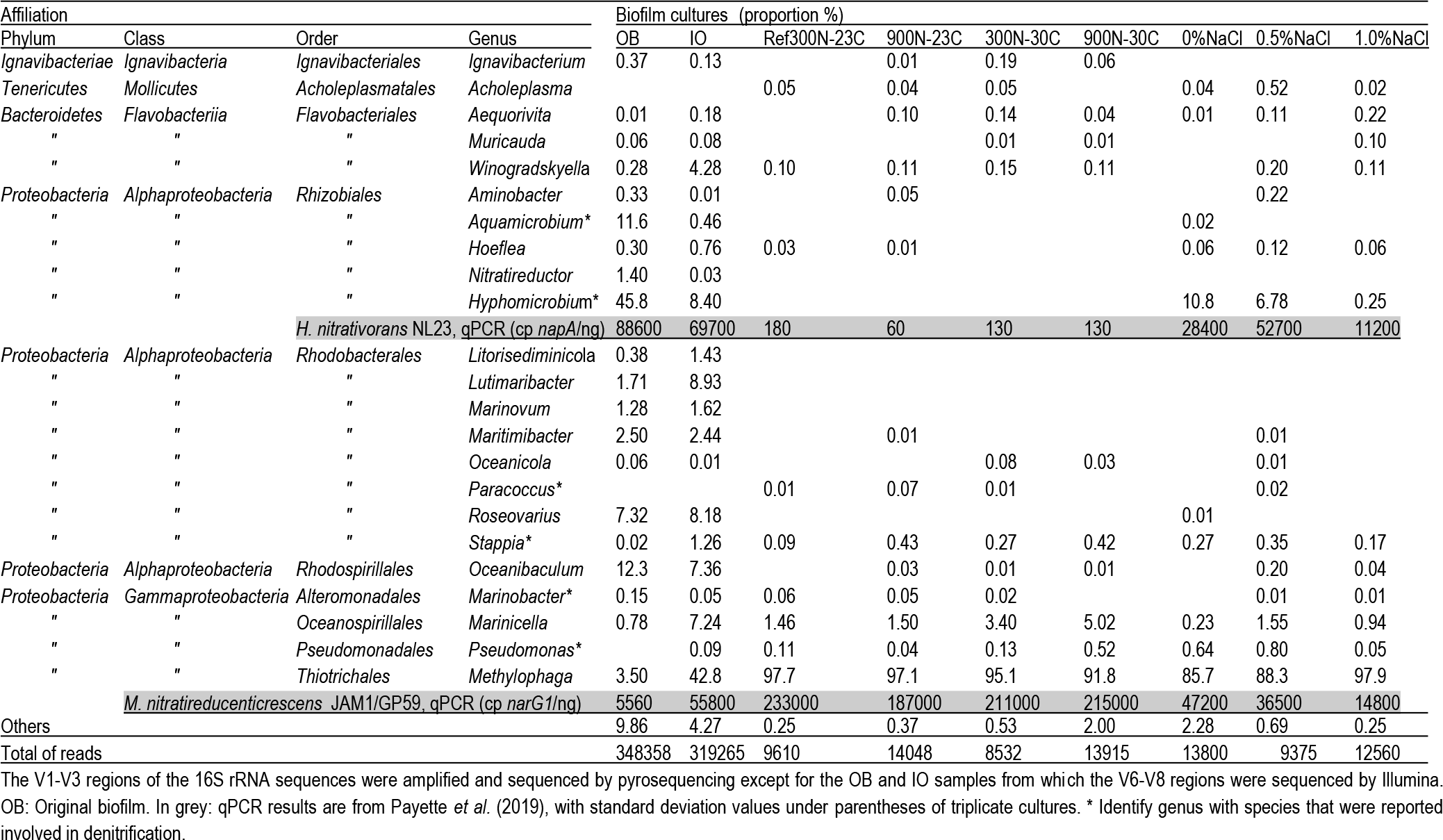
Most probable affiliation of the most abundant 16S rRNA sequences in the biofilm cultures and the original biofilm

In the OB, high proportions of the 16S sequences were related to *Hyphomicrobium* spp. (45.8%) followed by *Oceanibaculum* spp. (12.3%), *Aquamicrobium* spp. (11.6%); *Methylophaga* spp. accounted for 3.5% (Table 2). In the IO biofilm cultures, the proportion of *Methylophaga* spp. increased by *ca.* 12-fold (42.8%) compared to the OB, and decreased by 5.5-fold for *Hyphomicrobium* spp. (8.4%). Higher proportions of *Marinicella* spp. (7.2%) and *Winogradskyella* spp. (4.3%) along with a substantial decrease in proportion of *Aquamicrobium* spp. (0.44%) were observed in the IO biofilm cultures compared to the OB (Table 2).

In the biofilm cultures acclimated to the four combinations of NO_3_^−^ concentrations and temperatures in ASW medium containing 2.75% NaCl (Ref300N-23C, 300N-30C, 900N-23C, 900N-30C), *Methylophaga* spp. accounted for >90% of the 16S sequences followed by *Marinicella* spp. with proportions ranging from 1.5% to 5.0% (Fig. 2; Table 2). No sequences were found affiliated to *Hyphomicrobium* spp. under these conditions. In the biofilm acclimated to low NaCl concentrations (0%NaCl, 0.5%NaCl, 1%NaCl), *Hyphomicrobium* spp. accounted for 11.8%, 6.8% and 0.25%, respectively of the 16S sequences (Fig. 2; Table 2). *Methylophaga* spp. was still the dominant genus with more than 85% of the 16S sequences; *Marinicella* spp. 16S sequences were also found in significant proportions (Fig. 2; Table 2). 16S sequences affiliated to *Stappia* spp. were found in all examined biofilm cultures and in the OB.

The 16S sequences from the OB and the IO biofilm cultures that were derived by Illumina sequencing generated several thousands of reads affiliated to *Hyphomicrobium* spp. and *Methylophaga* spp. This tremendous amounts of sequences allowed assessing the presence of species other than *H. nitrativorans* and *M. nitratireducenticrescens* in these two biofilms. The phylogenic analyses performed on these sequences revealed three clusters of OTUs of *Hyphomicrobium* spp. and three clusters of *Methylophaga* spp. (Fig. 3A and 3B). The vast majority (>90%) of the 16S rRNA sequences associated to these OTUs were affiliated to *H. nitrativorans* or *M. nitratireducenticrescens*, respectively, in the OB and the IO biofilm cultures (Clusters 1, Table 3). A small proportion of the OTUs (clusters 2 and 3) was affiliated to other *Hyphomicrobium* or *Methylophaga*, which suggests that other members of these genera were present in these biomasses in very low proportions.

**Table 3:**
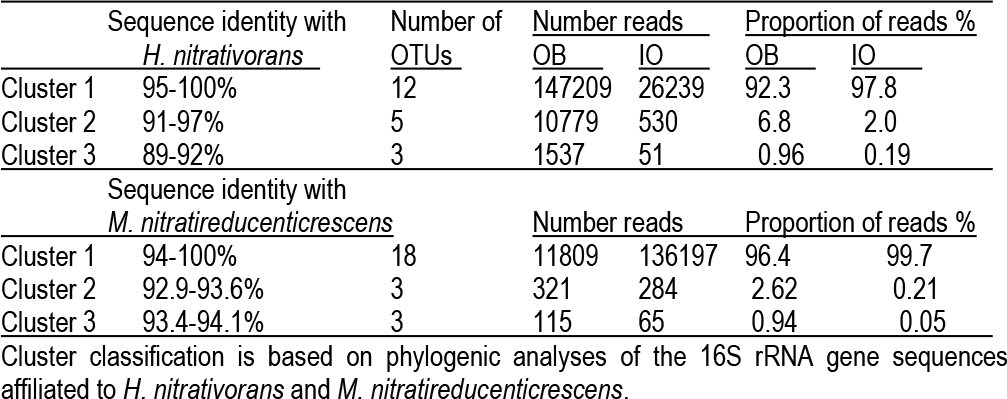
OTUs affiliated to *Hyphomicrobium* spp. and *Methylophaga* spp. in the OB and the IO biofilm cultures

**Figure 3.**
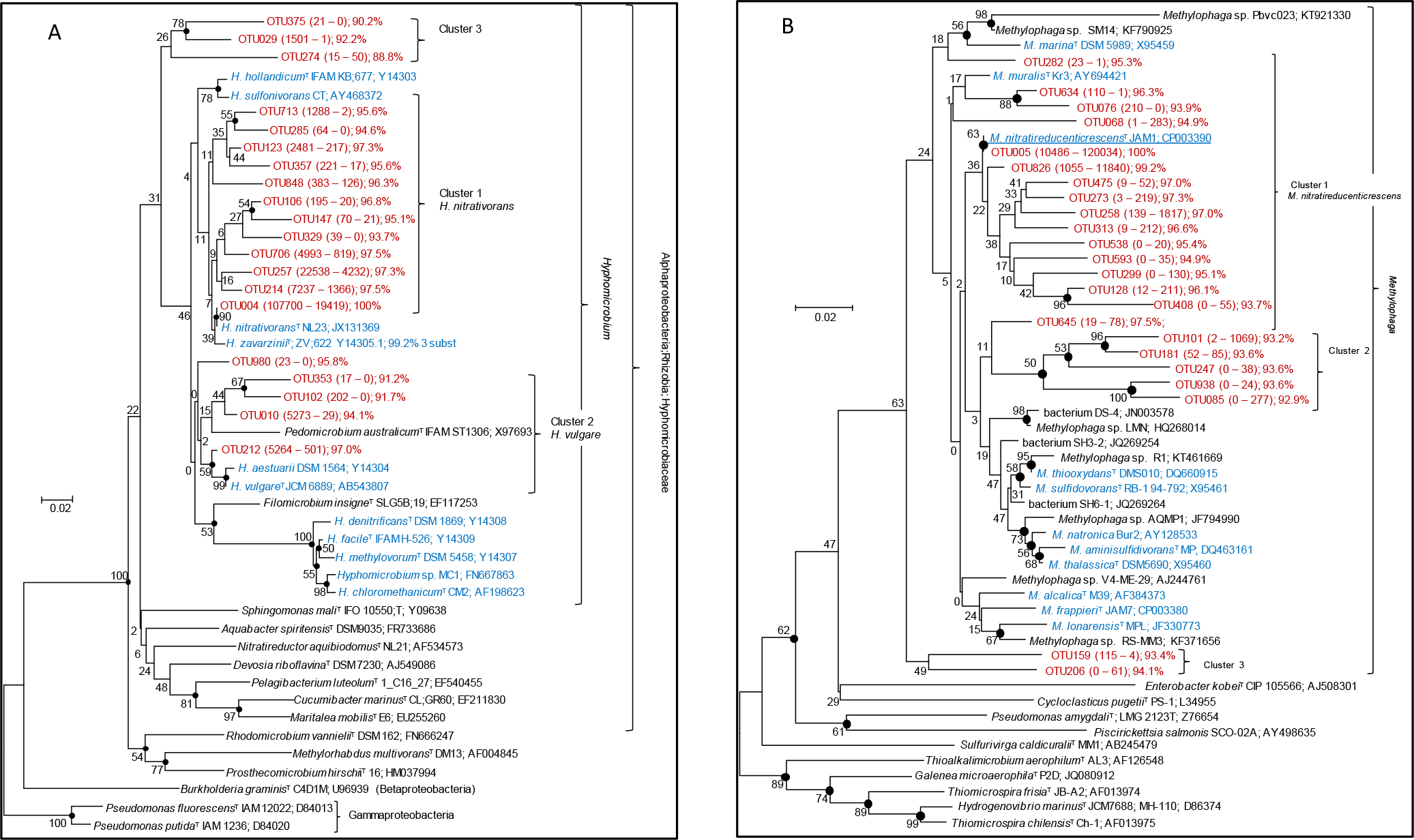
Evolutionary relationships of OTUs derived from the OB and the IO biofilm cultures with the genera *Hyphomicrobium* and *Methylophaga*. An unrooted phylogenetic tree demonstrating the evolutionary relationship of 16S rRNA gene sequences (V6-V7-V8 regions) is illustrated. The evolutionary history was inferred using the Minimum Evolution method. The percentage of replicate trees in which the associated taxa clustered together in the bootstrap test (1000 replicates) are shown next to the branches. The tree is drawn to scale, with branch lengths in the same units as those of the evolutionary distances used to infer the phylogenetic tree. The evolutionary distances were computed using the Maximum Composite Likelihood method and are in the units of the number of base substitutions per site. The rate variation among sites was modeled with a gamma distribution (shape parameter = 1.7). The ME tree was searched using the Close;Neighbor;Interchange (CNI) algorithm at a search level of 1. The Neighbor;joining algorithmwas used to generate the initial tree. All positions with less than 95% site coverage were eliminated. That is, fewer than 5% alignment gaps, missing data, and ambiguous bases were allowed at any position. There were a total of 409 positions in the final dataset. Evolutionary analyses were conducted in MEGA6 [6]. Beside OTU names are the number of reads derived from the OB and the IO biofilm cultures, respectively, and the percentage of identity with *H. nitrativorans* NL23 16S sequence (Panel A) or with *M. nitratireducenticrescens* JAM1 16S sequence (Panel B). Beside the species names is the GenBank accession number. Panel A: *Hyphomicrobium* The optimal tree with the sum of branch length = 1.70056439 is shown. The analysis involved 46 nucleotide sequences. Panel B: *Methylophaga* The optimal tree with the sum of branch length = 1.56790826 is shown. The analysis involved 54 nucleotide sequences.

### Isolation of bacterial isolates from the biofilm cultures

The biomass of the Ref300N-23C biofilm cultures was dispersed on different nutrient agar plates to isolate other denitrifying bacteria. Several isolates affiliated to *M. nitratireducenticrescens* were recovered from the methylophaga 1403 agar medium, among which strain GP59 was obtained (Geoffroy *et al.*, 2018). Isolates affiliated to the genera *Marinobacter, Pseudomonas, Paracoccus, Roseovarius, Thalassobius*, *Winogradskyella, Aequorivita* and *Exiguobacterium* (Table 4) were recovered from the Marine medium 2216 plates, which contains yeast extract and peptone (Atlas, 1993). Only isolates affiliated to the genera *Marinobacter* and *Paracoccus* showed consumption of NO_3_^−^ and NO_2_^−^ and production of gas, suggesting that they possess the complete denitrification pathway. The three isolates affiliated to the *Paracoccus* spp. have identical 16S rRNA sequences to each other and to the one of *Paracoccus* sp. strain NL8 that was isolated from the Biodome denitrification system (Labbé *et al.*, 2003). One representative of *Paracoccus* isolates (GP3) could grow with methanol as sole source of carbon; *Marinobacter* sp. GP2 could not.

**Table 4.**
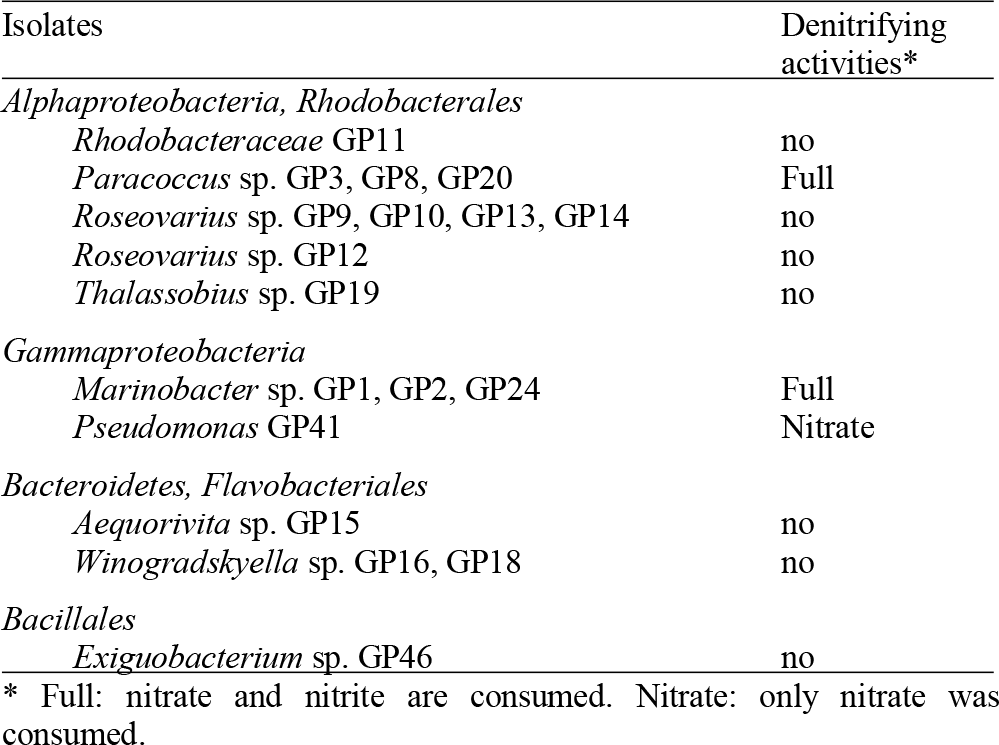
Affiliation of the isolates isolated from the Reference biofilm cultures (Ref300N-23C)

| Isolates | Denitrifying activities* |
| --- | --- |
| *Alphaproteobacteria, Rhodobacterales* |  |
| *Rhodobacteraceae* GP11 | no |
| *Paracoccus* sp. GP3, GP8, GP20 | Full |
| *Roseovarius* sp. GP9, GP10, GP13, GP14 | no |
| *Roseovarius* sp. GP12 | no |
| *Thalassobius* sp. GP19 | no |
| *Gammaproteobacteria* |  |
| *Marinobacter* sp. GP1, GP2, GP24 | Full |
| *Pseudomonas* GP41 | Nitrate |
| *Bacteroidetes, Flavobacteriales* |  |
| *Aequorivita* sp. GP15 | no |
| *Winogradskyella* sp. GP16, GP18 | no |
| *Bacillales* |  |
| *Exiguobacterium* sp. GP46 | no |

\* Full: nitrate and nitrite are consumed. Nitrate: only nitrate was consumed.

### Metatranscriptomic analysis of the biofilm cultures

The metatranscriptomic approach has allowed assessing the potential metabolic contributions of the microbial community in the biofilm cultures. We have chosen to focus on three biofilm cultures, which were the Ref300N-23C (the Reference biofilm cultures), 900N-30C (highest denitrification rates; Table 1) and 0%NaCl (persistence of *H. nitrativorans* NL23) biofilm cultures. Because the genomes of *H. nitrativorans* NL23 and *M. nitratireducenticrescens* JAM1 and GP59 were available, we first determined changes in the transcript levels of genes associated to these genomes between the biofilm cultures and the planktonic pure cultures of the respective strains. To assess the contributions of other microorganisms in biofilm cultures, the reads from the metatranscriptomes that did not align with the three reference genomes were used to derive *de novo* assembled transcripts. These transcripts were annotated for function and bacterial affiliation.

### Gene expression profiles of *M. nitratireducenticrescens* GP59 in the biofilm cultures

Because >80% of the genomes of strains JAM1 and GP59 are identical, high proportions of reads from the biofilm metatranscriptomes can align to both genomes. Geoffroy *et al.* (2018) showed that the gene expression profiles of the common genes between both strains in planktonic pure cultures were similar. Results from qPCR (Payette *et al.*, 2019) showed that the concentrations of strain GP59 were one to three orders of magnitude higher than those of strain JAM1 in the three biofilms cultures. Because of these differences, it was assumed that most of the transcript reads associated to *M. nitratireducenticrescens* in the biofilm cultures were from strain GP59. The transcriptomes of strain GP59 were also derived from planktonic pure cultures cultivated under anoxic conditions in the methylophaga 1403 medium (Fig. 1). The choice of this medium was because suboptimal growth occurred with strain GP59 in ASW medium. The relative transcript levels of the corresponding genes in the biofilm cultures and the planktonic pure cultures were compared to assess changes in the metabolisms of the strain that occurred between the two environments. All quantitative changes described below of the gene transcript levels in the biofilm cultures are expressed relative to the transcript levels in the planktonic pure cultures.

Among all GP59 genes, between 11% and 21% of them had higher relative transcript levels in the biofilm cultures. At the opposite, 6 to 17% of all GP59 genes were expressed at higher relative transcript levels in planktonic pure cultures (Fig. 4). Strain GP59 contains two plasmids, and most of the genes encoded by these plasmids had much lower relative transcript levels in the biofilm cultures (Fig. 4). Genes involved in the nitrogen metabolism and iron transport were globally at higher relative transcript levels in the biofilm cultures (Table 5; Fig. 4 and 5A).

**Figure 4.**
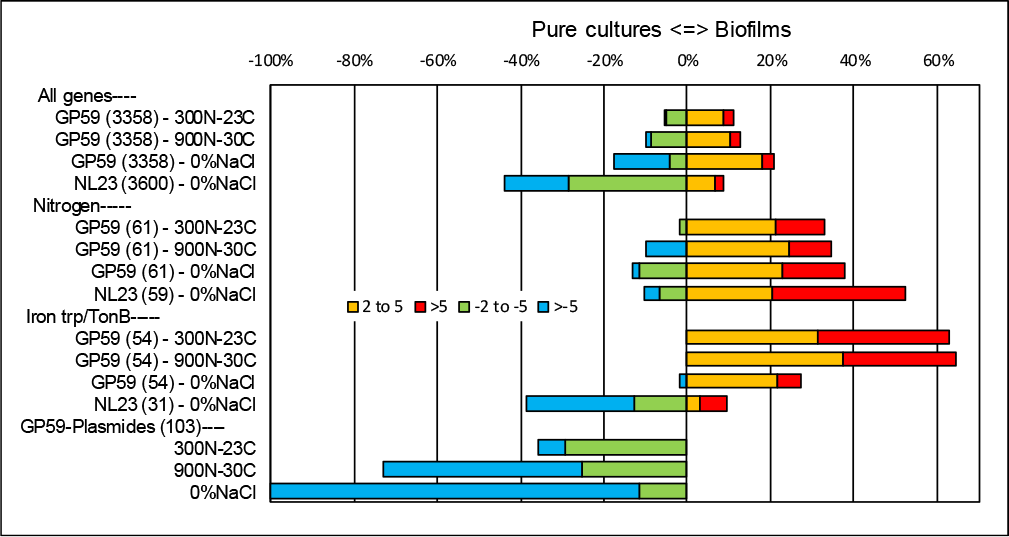
Relative expression profiles of *M. nitratireducenticrescens* GP59 and *H. nitrativorans* NL23 in biofilm cultures. All the deduced amino acid sequences associated to the GP59 genome and plasmids and the NL23 genome were submitted to the BlastKOALA (genome annotation and KEGG mapping) at the Kyoto encyclopedia of genes and genomes (KEGG). Genes associated to specific metabolisms were sorted out and the corresponding ratio of the Biofilm TPM versus the pure culture TPM was derived. When the ratios were <1, the negative inverse value (−1/ratio) was calculated. Data are expressed as the percentage of genes in each category that are more expressed in the biofilm cultures (right, 2 to 5 times, and > 5 times) or in pure cultures (left, −2 to −5 times and > −5 times). Number within parentheses are the number of genes involved in the selected pathways. Other metabolic profiles are detailed in Figures S1 and S2.

**Figure 5:**
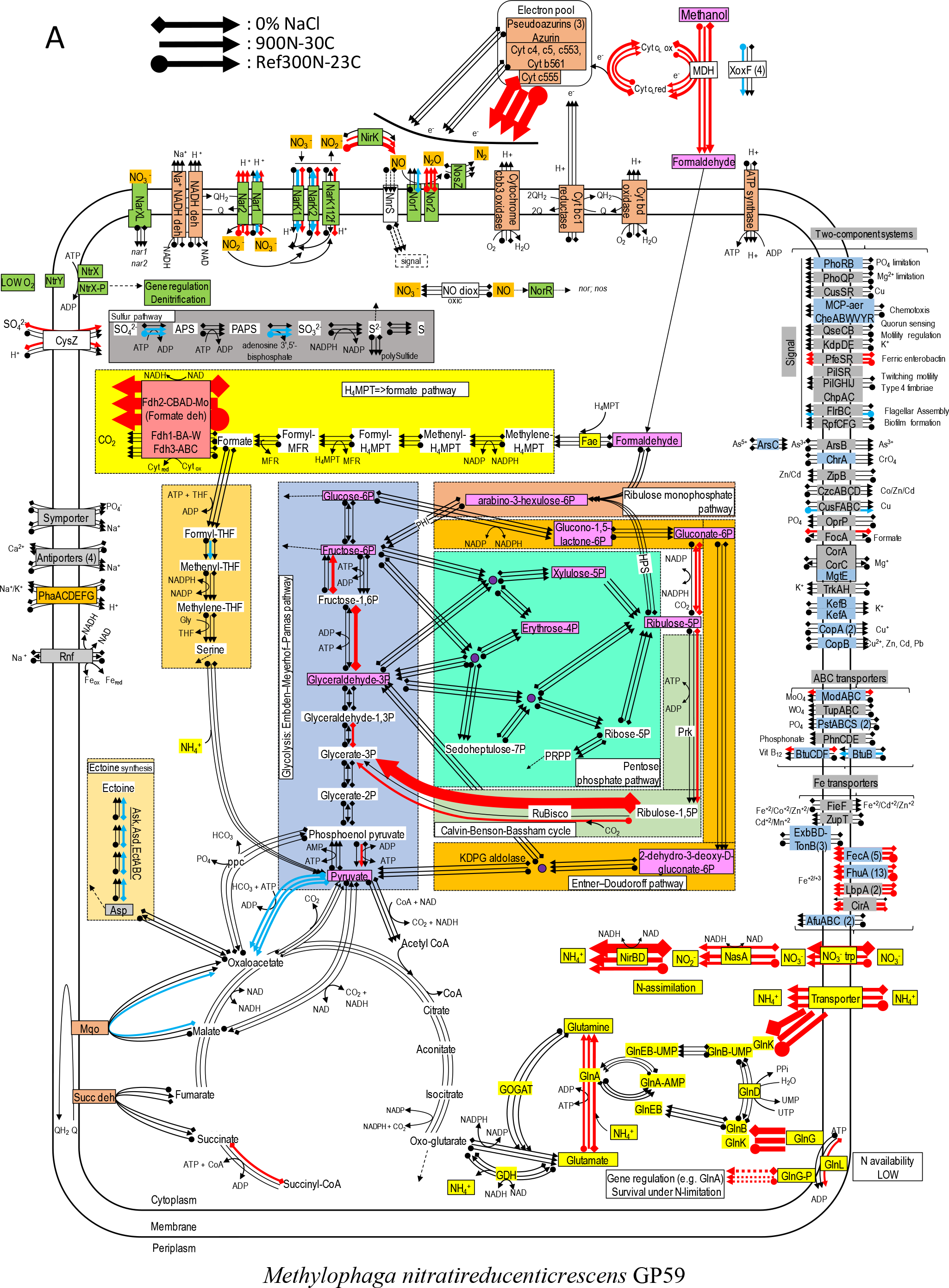
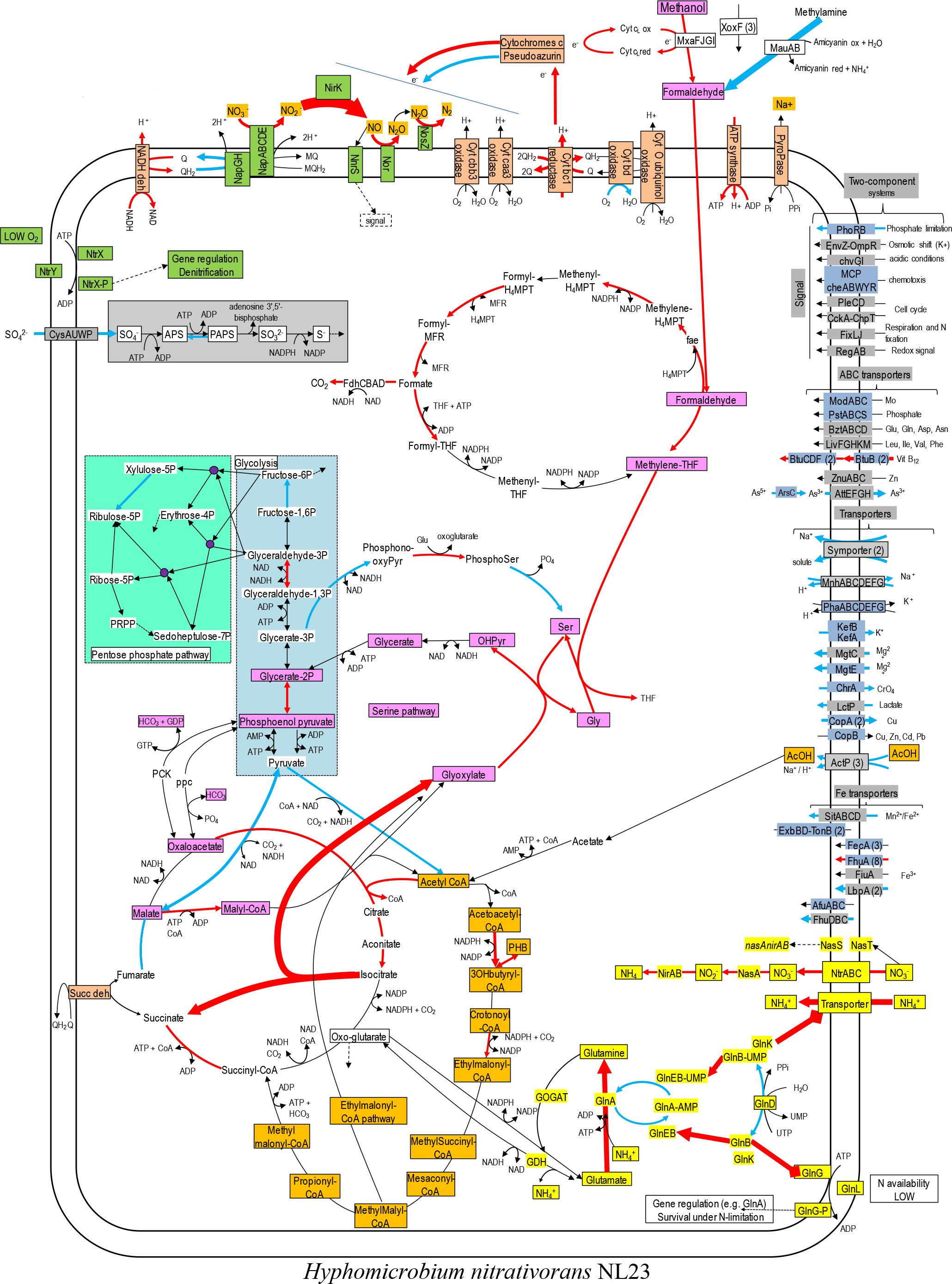
Relative gene expression profiles of selected metabolic pathways in *M. nitratireducenticrescens* GP59 and *H. nitrativorans* NL23 in the biofilm cultures. Panel A: *M. nitratireducenticrescens* GP59. Panel B: *H. nitrativorans* NL23. The pathways are based on functions deduced by the annotations (provided by KEGG BlastKoala, RAST and GenBank). The arrow thickness is proportional to the value of the ratio of the Biofilm TPM divided by the planktonic pure-culture TPM. The blue arrows represent genes with at least 2-fold lower relative transcript levels in the biofilm cultures. The red arrows represent genes with at least 2-fold higher relative transcript levels in the biofilm cultures. The black arrows represent no changes between both types of cultures in the relative transcript levels. The two-component systems and the transporters that are illustrated in blue are encoded by both strains.

**Table 5:**
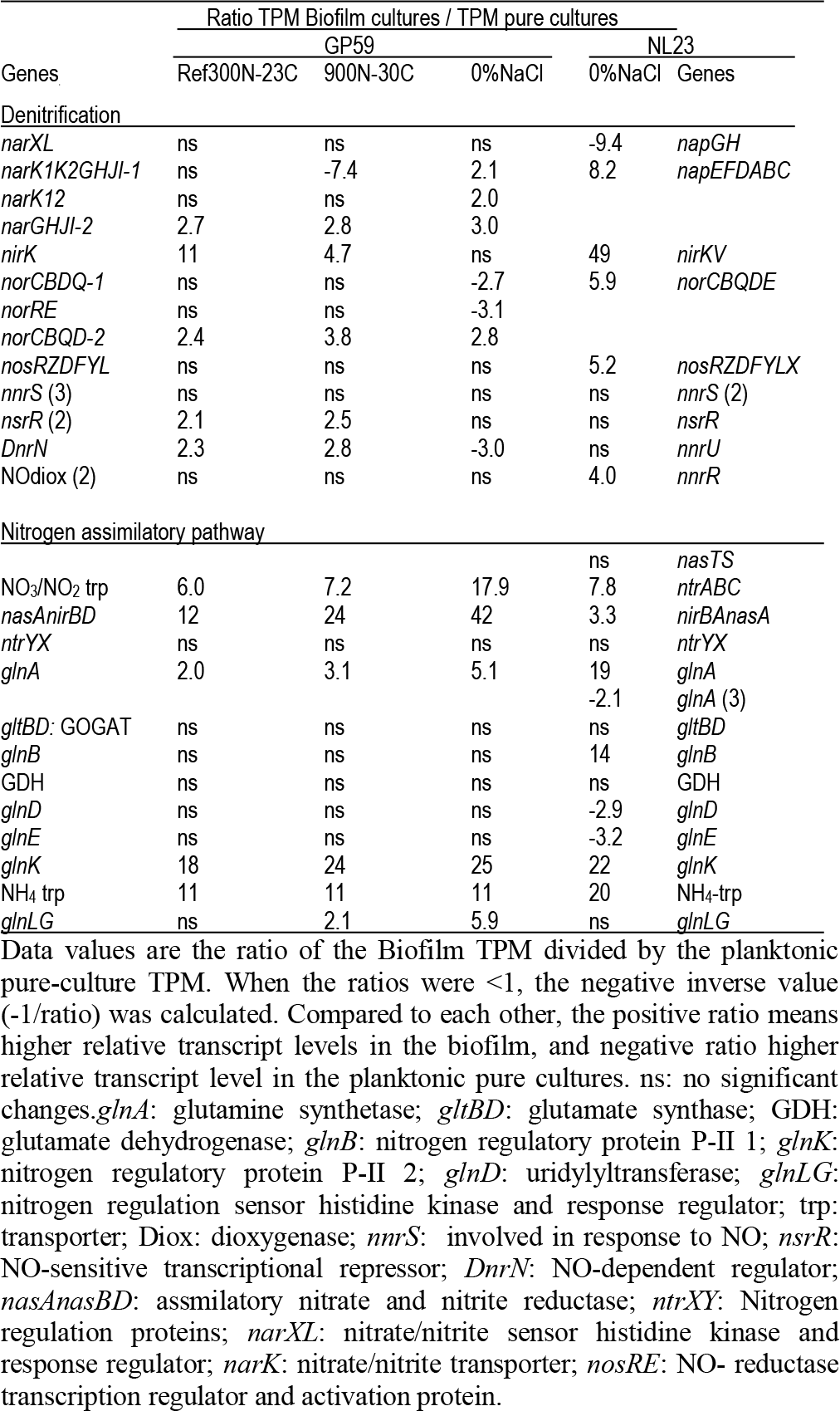
Changes in the relative transcript levels of genes involved in the nitrogen metabolism in strain GP59 and strain NL23

For the denitrification genes, *narXL* encoding the regulatory factors of the *nar* systems showed no differences between the biofilm cultures and the planktonic pure cultures in the relative transcript levels (Table 5). Small upregulation of the *nar2* operon with about 3-fold increases in relative transcript levels occurred in the biofilm cultures. These levels were much lower in the 900N-30C biofilm cultures for the *nar1* operon and were about the same levels in the two other biofilm cultures and the planktonic pure cultures. The *nor1* operon had the same relative transcript levels in the 300N-23C and 900N-30C biofilm cultures and the planktonic pure cultures, and a 3-fold decrease was noticed in these levels in the 0%NaCl biofilm cultures (Table 5). No significant changes in the expression of the *nos* operon occurred between the biofilm cultures and the planktonic pure cultures. The relative transcript levels of *nirK* were 5 to 10-times higher in the 300N-23C and 900N-30C biofilm cultures, whereas these levels were similar in the 0%NaCl biofilm cultures and the planktonic pure cultures (Table 5). Higher relative transcript levels of genes involved in the ammonium transport and the assimilatory NO_3_^−^/NO_2_^−^ reduction pathway were observed in the biofilm cultures (Table 5; Fig. 5A). Absence of nitrogen source other than NO_3_^−^ in the ASW medium and presence of 37 mM NH_4_^+^ in the medium used for the planktonic pure cultures (methylophaga 1403) could explain these differences in the assimilatory pathway.

Figure 5A illustrates changes in the relative expression profiles in the biofilm cultures of major pathways in strain GP59. The relative transcript levels of *mxaFJGI* encoding the small and large subunits of the methanol dehydrogenase (MDH) and the cytochrome c-L increased by 2-6 fold in biofilm cultures. Two out of the four *mxaF*-related products (*xoxF*) showed 2- to 9-fold decreases in their relative transcript levels in the biofilm cultures. As observed in *Methylorubrum extorquens*, the *M. nitratireducenticrescens* GP59 genome encodes three formate dehydrogenases with the same gene arrangement (Chistoserdova *et al.*, 2004). The *fdhCBAD* operon that encodes the NAD-linked, Mo-formate dehydrogenases showed *ca.* 60-fold increases in the relative transcript levels in the biofilm cultures, whereas the two other *fdh* operons stayed at the same levels of those of the planktonic pure cultures. The strong upregulation of this operon may be related to the increasing need of NADH in the biofilm. The relative transcript levels of the gene encoding the cytochrome c555 were 40 to 70 times higher in the biofilm cultures. Genes encoding the other cytochromes, pseudoazurins and azurin were expressed at similar levels in the biofilm cultures and the planktonic pure cultures. Genes involved in the formaldehyde metabolism to formate and CO_2_, glycolysis, the ribulose monophosphate pathway, the Entner Doudorof pathway, the tricarboxylic acid cycle, and the pentose pathway showed their relative transcript levels in general unchanged. Few genes in these pathways had 2-11 fold differences between the planktonic pure cultures and the biofilm cultures. The GP59 genome encodes the major enzymes involved in the Calvin-Benson-Bassham cycle: the ribulose-bisphosphate carboxylase (Rubisco) and the phosphoribulokinase (Prk). In the 0%NaCl biofilm cultures, the relative transcript levels of the Rubisco gene operon (*rbcSL*) jumped by 66 times. This upregulation was less pronounced in the Ref300N-23C biofilm cultures (3-fold increase). For the prk gene, the relative transcript levels were 3 to 4 times higher in the 0%NaCl biofilm cultures. The nature of this upregulation is unknown. Except for the cytochrome c555, the relative transcript levels of genes encoding for the oxidative phosphorylation metabolism were unchanged. Several genes involved in iron transport showed higher relative transcript levels in the biofilm cultures (2 to >50-fold increases). The nature of this upregulation in the biofilm cultures is unknown, as the biofilm and planktonic pure cultures were cultivated with trace elements containing iron (Payette *et al.*, 2019).

### Gene expression profiles of *H. nitrativorans* NL23 in the 0%NaCl biofilm cultures

As with *M. nitratireducenticrescens* GP59, the transcript levels of genes associated to *H. nitrativorans* NL23 were compared between the biofilm cultures and the planktonic pure cultures to assess changes in the metabolisms that occurred between the two environments. The NL23 planktonic pure cultures were cultivated under anoxic conditions in the 337a medium (Fig. 1). The choice of this medium was because strain NL23 could not grow in ASW (with 2.75% NaCl).

The overall analysis of the three metatranscriptomes confirmed the results obtained by qPCR assays and the 16S rRNA gene analysis (Table 2). High number of reads (40 × 10^6^) derived from the metatranscriptome of the 0%NaCl biofilm cultures aligned with the NL23 genome, but <20 000 reads derived from the Ref300N-23C and 900N-30C metatranscriptomes did. In the 0%NaCl biofilm cultures, <10% of all NL23 genes had a higher relative transcript levels in the 0%NaCl biofilm cultures, whereas this was the case for >40% genes in planktonic pure cultures (Fig. 4). These results suggest important changes had occurred in the regulation of gene expression between the planktonic pure cultures and the biofilm cultures. Genes involved in the energy (Fig. S2) and nitrogen metabolisms (Fig. 4; Table 5) had globally higher relative transcript levels in the 0%NaCl biofilm cultures.

Higher relative transcript levels (5 to 8-times) for the *nap*, *nor* and *nos* operons were observed in the 0%NaCl biofilm cultures (Table 5). *nirK* was highly upregulated in the biofilm cultures with 49-fold increase in the relative transcript levels (Table 5). The *napGH* operon however had a 9.4-fold decrease in the relative transcript levels in the 0%NaCl biofilm cultures. As observed with strain GP59, substantial changes in the relative transcript levels of genes involved in the ammonium transport and the assimilatory NO_3_^−^/NO_2_^−^ reductase were observed with 3- to 22-fold increases in these levels (Table 5, Fig. 5B). These results correlate with the absence of NH_4_^+^ in the ASW medium, and thus NO_3_^−^ the only source of N, compared to the 337a medium used for the planktonic pure cultures, which contains 3.8 mM NH_4_^+^.

Figure 5B illustrates changes in relative expression profiles of major pathways in the 0%NaCl biofilm cultures of strain NL23. The relative transcript levels of *mxaFJGI* increased by 2-fold in the biofilm cultures. The relative transcript levels of the *mau* operon (methylamine dehydrogenase) showed a 15-fold decrease in the biofilm cultures. The nature of such decrease is unknown as strain NL23 was not fed with methylamine in any of our cultures. The three *xoxF* genes did not show substantial changes in their transcript levels in both types of cultures. Genes involved in the formaldehyde metabolism to formate and CO_2_, glycolysis, the tricarboxylic acid cycle, and the pentose pathway showed their transcript levels in general unchanged between the planktonic pure cultures and the biofilm cultures. Few genes in these pathways had 2-5 fold differences in their relative transcript levels The two genes encoding the key enzymes in the serine pathway (alanine-glyoxylate transaminase, glycine hydroxymethyltransferase) had 3- to 7-fold increases in their relative transcript levels in the biofilm cultures. As *Methylorubrum extorquens*, the NL23 genome encodes the ethymalonyl-CoA pathway (Chistoserdova *et al.*, 2003; Peyraud *et al.*, 2009), which did not show changes overall in the transcript levels of the corresponding genes between the two types of cultures. Contrary to *M. extorquens* however, a gene encoding the isocitrate lyase is present in strain NL23 and showed a 25-fold upregulation in the biofilm cultures. The isocitrate lyase is one of the key enzymes of the glyoxylate bypass that catalyzes the transformation of isocitrate to succinate and glyoxylate. Gene encoding isocitrate lyase is also present in other available *Hyphomicrobium* genomes. All these results suggest that in the 0%NaCl biofilm cultures, the carbon metabolism increased in activity and that the glycine regeneration for the serine pathway by the glyoxylate was upregulated. Among genes involved in the oxidative phosphorylation, the relative transcript levels were higher (2 to 13 times) in the biofilm cultures with those encoding the NADH dehydrogenase, the cytochrome c reductase, with one of the cytochromes c and the F-type ATPase. Combined with increases in the relative transcript levels of the denitrification and the carbon pathways, these results suggest that increases in electron donor activities correlates with the need of electron for the nitrogen dissimilatory metabolism in the biofilm cultures. Strain NL23 possesses four types of cytochrome oxidase (aa3, bo, bd-I and cbb3) (reduction of O_2_ in H_2_O) that are in general expressed at the same levels in the biofilm and the planktonic pure cultures. Figure 5B also illustrates the dynamic changes of transporters and two component systems. Several of these transporters had a lower relative transcript levels in the 0%NaCl biofilm cultures. Contrary to strain GP59, genes involved in iron transport were not strongly affected in their gene expression in the 0%NaCl biofilm cultures (Fig. 4 and 5B).

### The composition of the active microbial community in the biofilm cultures

As mentioned above, the reads from the three metatranscriptomes that did not align with the genomes of *H. nitrativorans* NL23 and *M. nitratireducenticrescens* GP59 and JAM1 were *de novo* assembled. These reads were subsequently aligned to the *de novo* assembled transcripts to derive the relative levels of these transcripts in the biofilm cultures. The *de novo* assembled sequences were then annotated for function and affiliation. Finally, these sequences were grouped by microbial affiliation to determine the active populations in the biofilm cultures and to assess their level of involvement in these biofilm cultures (Table 6).

It was estimated that between 5 to 10% reads of the three metatranscriptomes were derived from other microorganisms than *H. nitrativorans* NL23 and *M. nitratireducenticrescens* GP59 and JAM1. The proportions of transcripts having genes with no affiliation and transcripts with no genes represented between 31 and 53% of the *de novo* assembled transcripts. The proportions of transcripts affiliated to Archaea and Eukarya accounted together for <0.1% (Table 6), which suggests very low abundance of these microorganisms in the biofilm cultures. The proportions of transcripts affiliated to viruses, phages and plasmids in the *de novo* assembled transcripts represented between 0.6 and 21.7% (Table 6).

**Table 6.**
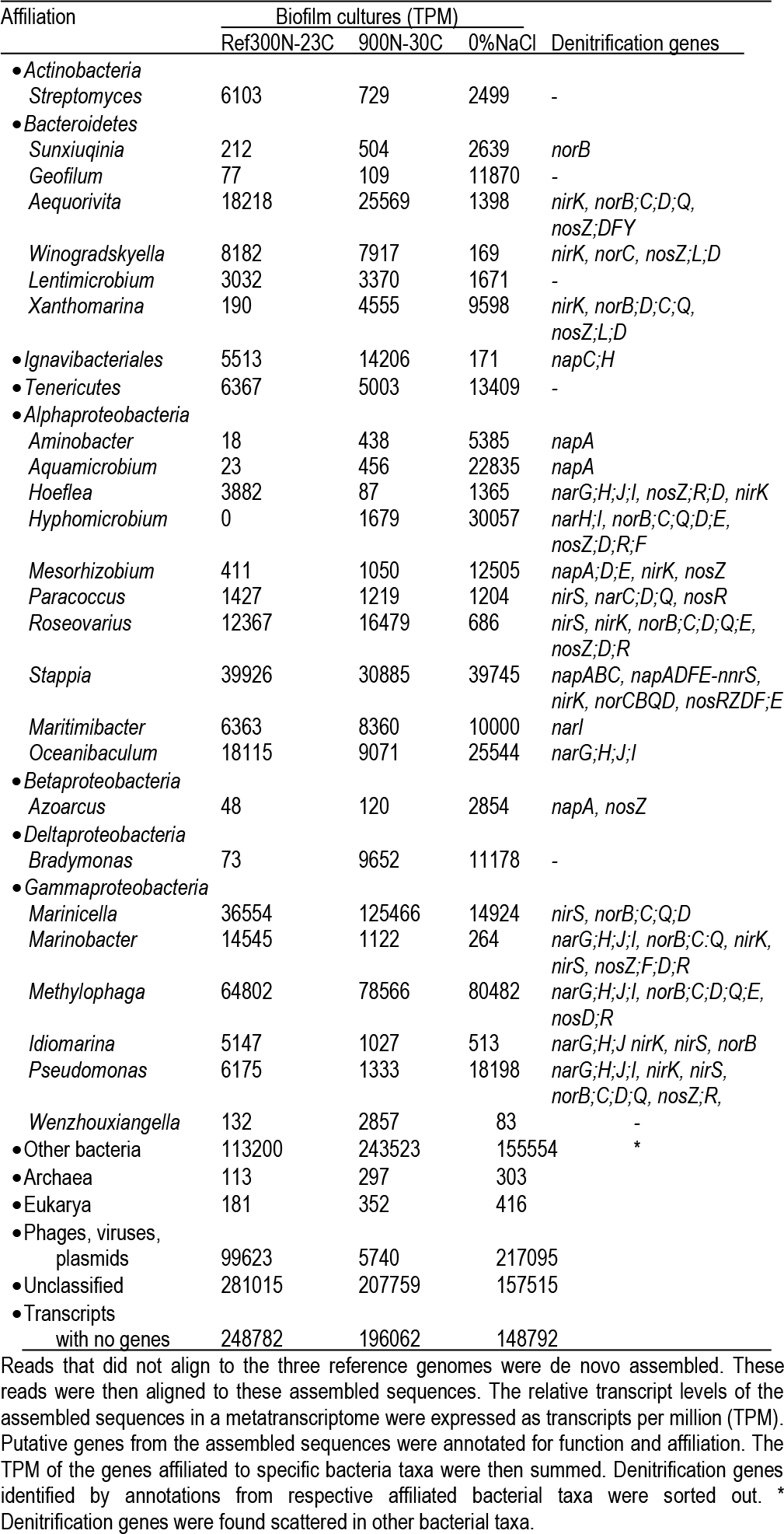
Microbial diversity and the associated denitrification genes in the biofilm cultures from the *de novo* transcript assembly

Twenty-seven bacterial taxa were selected for their overall transcript levels in at least one of the three biofilm cultures (Table 6). All the taxa detected by the 16S metagenomic approach are present in this list (Table 2). These 27 taxa represented between 22% and 35% of the *de novo* assembled transcripts. Among these taxa, genes encoding the four denitrification reductases were present in the *de novo* transcripts affiliated to *Marinobacter* spp., *Stappia* spp. and *Pseudomonas* spp. However, only *de novo* assembled transcripts affiliated to the *Stappia* spp. showed the complete set of denitrification genes in the three biofilm cultures and organized in operons (*napABC*, *napADFE*, *norCBQD*, *nosRZDF)*.

The expression profiles of the 27 bacterial taxa were compared between each biofilm culture by clustering analysis to assess whether some taxa were influenced in their global metabolic activities by the specific conditions of the biofilm cultures (Fig. 6). NaCl concentration was the main factor of clustering as two distinct clusters were derived. The low salt cluster consisting of nine bacterial taxa showed higher relative transcript levels in the 0%NaCl biofilm cultures, whereas the marine cluster of 13 bacterial taxa had higher relative transcript levels in the Ref300N-23C and 900-30C biofilms cultures. A third cluster showed five bacterial taxa with lower relative transcript levels in the 900N-30C biofilm cultures compared to the 0%NaCl and Ref300N-23C biofilm cultures. A third cluster showed five bacterial taxa with lower relative transcript levels in 900N-30C biofilm cultures compared to the 0%NaCl and Ref300N-23C biofilm cultures. In these cases, higher temperature (30°C vs 23°C) and higher NO_3_^−^ concentration (64.3 mM vs 21.4 mM) may have negatively affected these populations.

**Figure 6.**
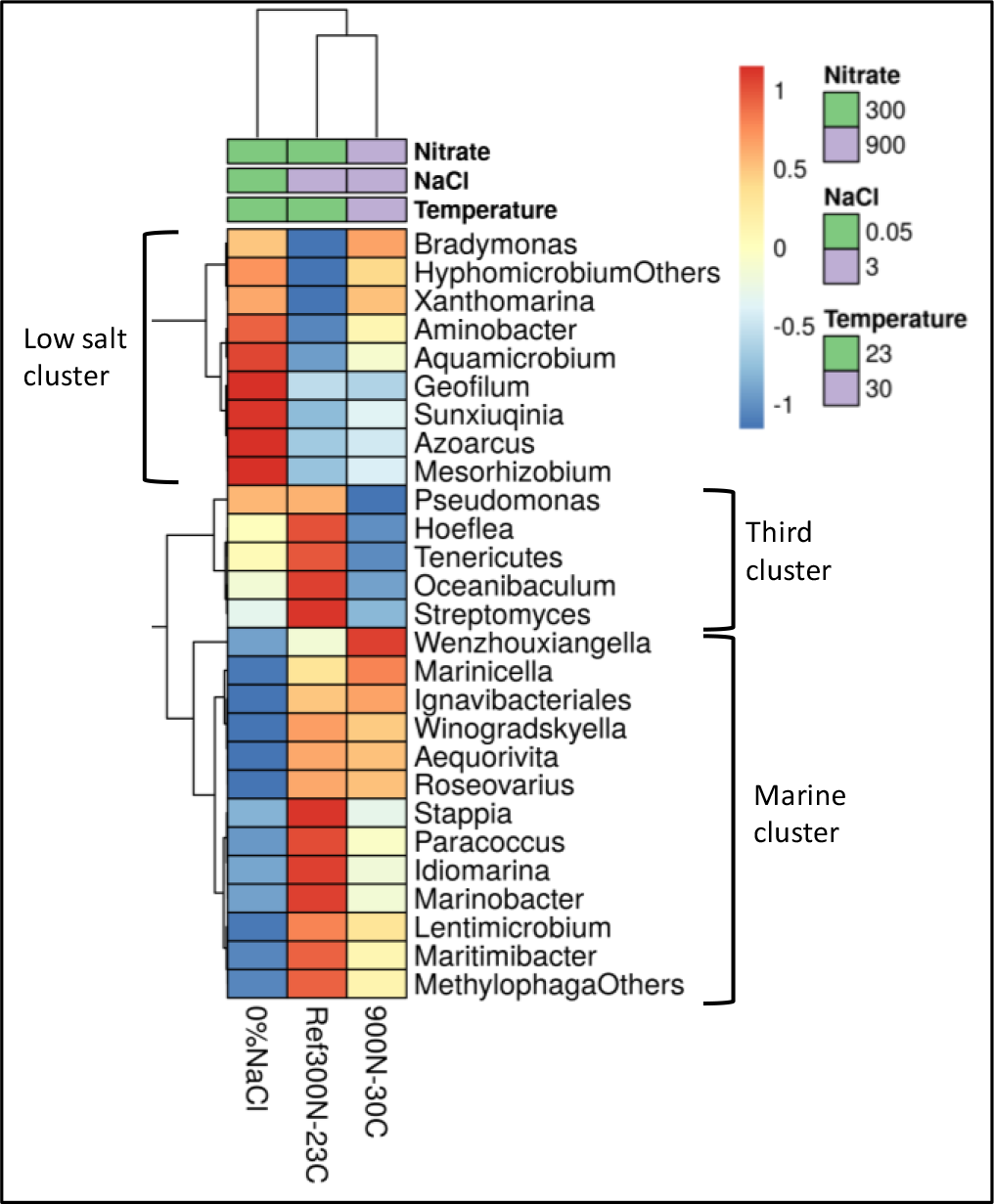
Hierarchical clustering of the selected bacterial taxa in the biofilm culture metatranscriptomes. Heatmap represents differences in the TPM (from Table 6) (log_10_ [TPM by geometric average of TPM]) between the three biofilm cultures for the respective bacterial taxa. Analysis was performed at ClustVis web site (https://biit.cs.ut.ee/clustvis/) (Metsalu and Vilo, 2015).

## Discussion

In the environment, numerous bacteria belonging to different taxa can accomplish denitrifying activities, and many of them were encountered in different types of denitrification processes (Lu *et al.*, 2014). Very few studies describing the microbial community of methanol-fed denitrification systems have been reported so far. Most of these studies are based on cloned 16S rRNA gene sequences of around 100 clones or based on fluorescence *in situ* hybridization (Neef *et al.*, 1996; Ginige *et al.*, 2004; Hallin *et al.*, 2006; Osaka *et al.*, 2006; Yoshie *et al.*, 2006; Baytshtok *et al.*, 2008; Osaka *et al.*, 2008; Rissanen *et al.*, 2016; Sun *et al.*, 2016; Rissanen *et al.*, 2017). In all these studies, high proportions of *Hyphomicrobium* spp. were found in combination with high proportions of other methylotrophs such as *Methyloversatilis* spp., *Methylophilus* spp., *Methylotenera* spp. or *Paracoccus* spp. The Biodome marine denitrification system showed no exception to this trend with co-occurrence of *Hyphomicrobium* spp. and the marine methylotroph *Methylophaga* spp. This co-occurrence was also observed in two other denitrification systems treating saline effluents (Osaka *et al.*, 2006; Rissanen *et al.*, 2016) (see Discussion by Payette *et al.*, 2019). The bacterial diversity of the biofilm taken from Biodome denitrification system was assessed before when the reactor was sampled in 2002 by deriving a 16S rRNA gene library and by culture approach (Labbé *et al.*, 2003; Labbé *et al.*, 2004). Beside *Hyphomicrobium* sp. and *Methylophaga* sp., *Paracoccus* sp., *Sulfitobacter* sp., *Nitratireductor aquibiodomus*, and *Delftia* sp. among others were identified. In the present report, a more complete determination of the composition of the bacterial community of the denitrifying biofilm that was frozen in 2006 (when the denitrification system was dismantle by the Biodome) was possible with the 16S metagenomic approach, as the following of the evolution of this community in the biofilm cultures. The metatranscriptomic approach provided a complementary method to determine this composition, but also provided good indications of the active metabolisms of the corresponding populations in the biofilm cultures. Our study is the first that gave a comprehensive picture of the microbial ecology and its evolution in a methylotrophic denitrifying biofilm.

The 16S metagenomic data from the OB and the IO biofilm cultures were derived from a different region of the 16S rRNA genes (V6-V8) than that used for the other biofilm cultures (V1-V3), and sequenced by a different technology (Illumina vs. pyrosequencing). The two approaches generated consistent results with qPCR (Table 2) that targeted *narG1* for *M. nitratireducenticrescens* and *napA* for *H. nitrativorans* NL23 (Payette *et al.*, 2019). In the biofilm cultures acclimated in ASW medium with 2.75% NaCl (Ref300N-23C, 300N-30C, 900N-23C and 900N-30C), *H. nitrativorans* NL23 concentrations determined by qPCR was close to the limit of detection, which concurs with the absence of 16S sequences affiliated to *Hyphomicrobium* spp. in these biofilm cultures. The 16S metagenomic data also showed, as qPCR, the persistence of strain NL23 in biofilm cultures acclimated in ASW medium with low NaCl concentrations (0%NaCl, 0.5%NaCl and 1.0%NaCl biofilm cultures). Finally, the concentrations of *H. nitrativorans* NL23 measured by qPCR were at the same levels in the OB and the IO biofilm cultures. The lower relative proportion of 16S sequences of *Hyphomicrobium* spp. in the IO biofilm cultures compared to OB was a consequence of the substantial growth of *M. nitratireducenticrescens* in these cultures, with a 10-fold increase in concentration as revealed by qPCR (Table 2). Grob *et al.* (2015) also observed strong growth of *Methylophaga* spp. in their seawater samples that were fed with 100 µM methanol with relative proportions of 16S sequences raising from <0.5% at T=0 to 84% after 3 days.

Results from the 16S metagenomes showed that *Marinicella* spp. were present in the OB and all the biofilm cultures and were the second most abundant bacterial population in the biofilm cultures acclimated in ASW containing 2.75% NaCl. These results concur with the metatranscriptomes of the biofilm cultures where *Marinicella* spp. had the relative transcript levels among the highest in the *de novo* assembled transcripts. *Marinicella* spp. are considered strict aerobic bacteria with no indication of NO_3_^−^ reduction (Romanenko *et al.*, 2010) although a previous study reported high relative abundances of *Marinicella* spp. in anoxic sulfide oxidizing reactors in which nitrate was used as the electron acceptor (Huang *et al.*, 2015). Genome annotations of two *Marinicella* strains (GenBank: *Marinicella* sp. F2 - assembly number ASM200005v1 and *M. litoralis* KMM 3900 - ASM259191v1) (Wang *et al.*, 2018) did not reveal complete denitrification pathway, beside a nitric oxide reductase gene cluster, also detected here in our metatranscriptomes. Together with the presence of *nirS* gene (Table 6), these results indicated that *Marinicella* spp. might have the capacity to use intermediates of the denitrification cycle to support their growth.

The 16S metagenomic data provided evidence of the presence of *Pseudomonas* spp., *Marinobacter* spp., *Stappia* spp., *Paracoccus* spp. and *Aquamicrobium* spp. in the OB and in the biofilm cultures. Some species belonging to these genera were reported to carry denitrification. Isolates affiliated to the genera *Pseudomonas*, *Marinobacter* and *Paracoccus* were recovered from the Ref300N-23C biofilm cultures. The *Marinobacter* and *Paracoccus* isolates could perform denitrifying activities and grow under anoxic conditions, whereas the *Pseudomonas* isolate could only consume NO_3_^−^.

Although denitrification genes were found in several of the bacterial populations identified by the metatranscriptomic approach, only transcripts encoding the four denitrification reductases affiliated to *Stappia* spp. were found in the three examined biofilm cultures. The proportions of *Stappia* spp. in the 16S metagenomes of these biofilm cultures ranged between 0.09% and 0.42% (Table 2). *Stappia* spp. are chemoorganotrophic bacteria found in marine environments (Weber and King, 2007) that can oxidize carbon monoxide (CO). They possess the form I *coxL* gene encoding the large subunit of CO dehydrogenase. Some also contain a gene for the large subunit of ribulose-1,5-bisphosphate carboxylase (RuBisCO) and may be able to couple CO utilization to CO_2_ fixation (King, 2003). A *coxL* gene was found in the *de novo* assembled transcripts affiliated to *Stappia* spp., but not RuBisCO. Sequences analogue to transporters for simple and multiple sugars such as xylose and fructose, and acetate were found, which suggest that the *Stappia* bacteria fed on the biofilm material for carbon sources. Combined with the isolation of denitrifying isolates affiliated to *Marinobacter* spp. and *Paracoccus* spp., these results suggest that the biofilm has the potential to adapt to heterotrophic non-methylotrophic environments.

The proportion of *Bacteroidetes* in the 16S metagenomes of the OB and the eight biofilm cultures ranged from 0.2% to 4.9%, and several genera of this phylum were identified in the three metatranscriptomes. Significant proportions of bacteria affiliated to the *Bacteroidetes* phylum were also found in other methanol-fed denitrification systems. For instance, 29% of cloned 16S rRNA sequences were affiliated to *Bacteroidetes* in an acclimated activated sludge in a methanol-fed anoxic denitrification process treating a synthetic wastewater with 4% NaCl (Osaka *et al.*, 2008). Isolates affiliated to the *Bacteroidetes Aequorivita* spp. and *Winogradskyella* spp. were isolated from the Ref300N-23C biofilm cultures. None of these two isolates, however, could sustain growth under denitrifying conditions. Although denitrification genes affiliated to *Bacteroidetes* genera were found in *de novo* assembled transcripts, genes encoding all the four denitrification reductases were not found to any of them. These results suggest that *Bacteroidetes* are not involved in denitrification, although they may be involved in some steps of the denitrification pathway.

The metatranscriptomic data provided some insights of specific metabolisms in *H. nitrativorans* NL23 and *M. nitratireducenticrescens* GP59 that were regulated in the biofilm environment. In absence of *H. nitrativorans* NL23 in the Ref300C-23C and the 900N-30C biofilm cultures, the GP59 *nirK* gene was upregulated by 5 to 10 times compared to the planktonic pure cultures that was also cultivated under denitrifying conditions. On the contrary, the relative transcript levels of this gene did not change between the 0%NaCl biofilm cultures and the planktonic pure cultures, while the relative transcript levels of the NL23 *nirK* were 49 times higher in the 0%NaCl biofilm cultures. These results suggest coordination in the expression of *nirK* between these two strains in the 0%NaCl biofilm cultures.

The gene clusters encoding the three other denitrification reductases (*nap, nor, nos*) in strain NL23 showed higher relative transcript levels in the 0%NaCl biofilm cultures. *napGH* was however down regulated in these biofilm cultures. *napGH* is located in a separate chromosomic region than the *napABCDEF* operon. NapGH and NapC have redundant function of transferring electrons to NapB across the membrane. It was proposed that NapC transfers electrons from the menaquinol, whereas NapGH do it from ubiquinol (Simon, 2011). The physiological consequence of *napGH* transcript decrease in the biofilm is unknown. Observations on *napEDABC* found in the denitrifier *Shewanella denitrificans* OS217, and *napDAGHB* in the respiratory NO_3_^−^ ammonifier *Shewanella oneidensis* MR-1 suggest that NapGH is more involved in the ammonification system (Simpson *et al.*, 2010). Despite the denitrifying conditions applied in both types of cultures, the biofilm environment has induced strong up regulation of denitrification genes in *H. nitrativorans* NL23. This may be in response to the rapid processing of NO_3_^−^ by *M. nitratireducenticrescens* JAM1/GP59 (Mauffrey *et al.*, 2015) that could generate rapidly high level of NO_2_^−^, which is toxic for strain NL23.

## Conclusion

This report concludes our study on the dynamics of the methylotrophic marine denitrifying biofilm originated from the denitrification system at the Montreal Biodome. We showed that the biofilm could be revived from years of frozen state in glycerol. The mature biofilm can sustain denitrifying activities with different NaCl concentrations, pHs and temperatures. Acclimation to ASW medium instead of the commercial IO medium that was used by the Biodome had a strong impact on the *Hyphomicrobium* population by the displacement of *H. nitrativorans* NL23 as the main denitrifier by a subpopulation of *M. nitratireducenticrescens*, represented by strain GP59 with the full denitrification pathway. However, low NaCl concentrations play an important role in the persistence of *H. nitrativorans* NL23 in the biofilm. In addition, for unknown reasons, this persistence occurred in the biofilm cultures acclimated in the IO medium, despite a NaCl concentration as high as found in the ASW medium. The biofilm environment has favored the upregulation of the denitrification pathway in *M. nitratireducenticrescens* GP59 and *H. nitrativorans* NL23 compared to planktonic pure cultures, despite the facts that these two types of cultures were grown under denitrifying conditions. Other denitrifiers affiliated to *Marinobacter* spp. and *Paracoccus* spp. were isolated from the biofilm cultures, and (an) uncultured denitrifier(s) affiliated to *Stappia* spp. was (were) metabolically active in the biofilm cultures. All these results demonstrated the dynamics and the plasticity of the denitrifying biofilm to sustain different environmental conditions and illustrate a comprehensive picture of the microbial community of the biofilm and its evolution during these changes. This could benefit in the development of optimal denitrifying bioprocess under marine conditions.

## ACKNOWLEDGEMENTS

We thank Karla Vasquez for her technical assistance.

## Funding

This research was supported by a grant to Richard Villemur from the Natural Sciences and Engineering Research Council of Canada # RGPIN-2016-06061. The funders had no role in study design, data collection and analysis, decision to publish, or preparation of the manuscript.

---

**Figure S1.**
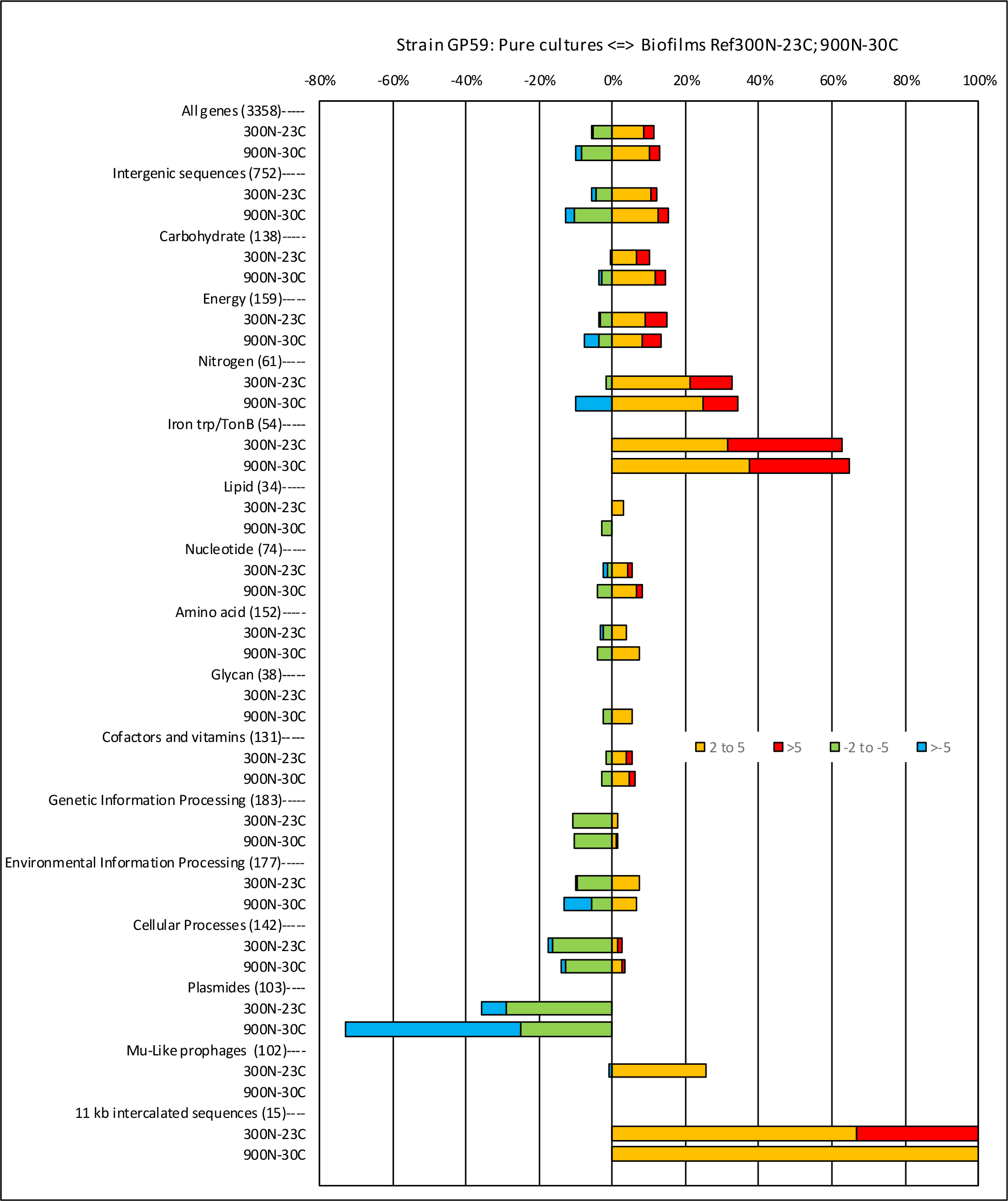
Relative expression profiles of *M. nitratireducenticrescens* GP59 in Ref300N-23C and 900N-30C biofilm cultures. All the deduced amino acid sequences associated to the GP59 genome and plasmids in the Ref300N-23C and 900N-30C biofilm cultures were submitted to the BlastKOALA (genome annotation and KEGG mapping) at the Kyoto encyclopedia of genes and genomes (KEGG). Genes associated to specific metabolisms were sorted out and the corresponding ratio of the Biofilm TPM versus the pure culture TPM was derived. When the ratios were <1, the negative inverse value (−1/ratio) was calculated. Data are expressed as the percentage of genes in each category that are more expressed in the biofilm cultures (right, 2 to 5 times, and > 5 times) or in pure cultures (left, −2 to −5 times and > −5 times). Number within parentheses is the number of genes involved in the selected pathways.

**Figure S2.**
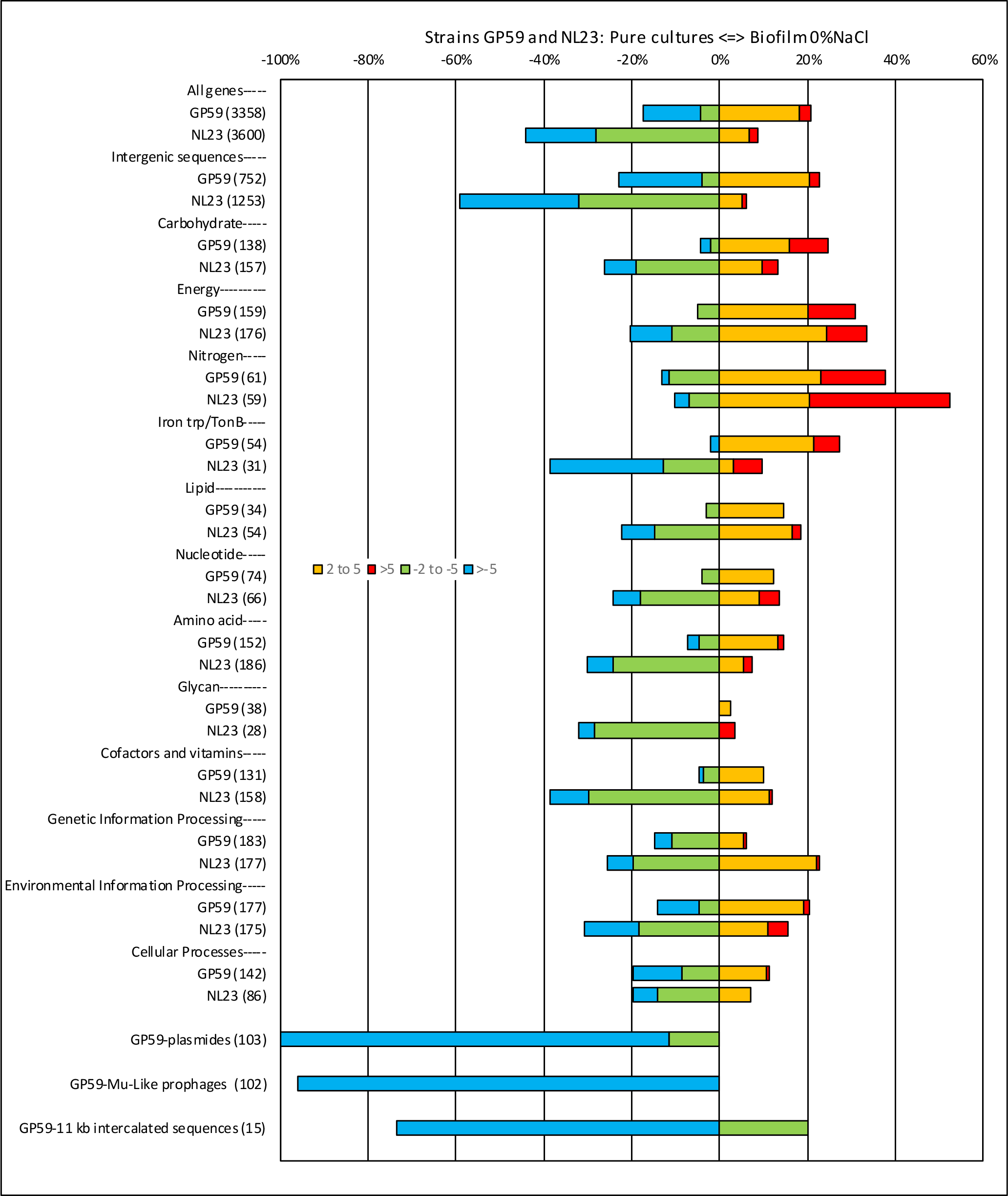
Relative expression profiles of *M. nitratireducenticrescens* GP59 and *H. nitrativorans* Nl23 in the 0%NaCl biofilm cultures. All the deduced amino acid sequences associated to the GP59 genome and plasmids and the NL23 genome in the 0%NaCl biofilm cultures were analyzed as described in Figure legend S1.

